# Transgenerational effects of heat shock on gene regulation and fitness-related traits are stronger in arid than temperate *Drosophila* populations

**DOI:** 10.1101/2025.02.13.637908

**Authors:** Ewan Harney, Josefa Gonzalez

## Abstract

Heat stress will increasingly affect populations as climate change leads to higher temperatures and more extreme events such as heat waves. Adaptation is predicted to occur over long evolutionary timescales, but recent work suggests that interactions between the epigenome and transposable elements (TEs) could link environmental acclimation with rapid adaptation. Yet combining all these factors in a tractable model system remains challenging; consequently little is known about how these processes interact in natural genetic backgrounds or shape evolutionarily relevant phenotypes.

To investigate these interactions, we carried out laboratory experiments measuring gene expression and chromatin accessibility responses to heat shock in female *D. melanogaster* from arid and temperate climates and their associations with population variation in TEs. The consequences for fitness-related phenotypes were characterised in the offspring, and we also measured expression, chromatin accessibility and phenotypic traits three generations later to explore transgenerational inheritance.

Expression and accessibility responses to heat shock varied between populations and were influenced by the presence of TEs, with more upregulated responses in the arid population. Effects of heat shock on transcription were detected three generations later, especially in the arid population, with many epigenetic genes transgenerationally expressed, although this was not driven by chromatin accessibility. Heat shock negatively affected phenotypes in the initial offspring cohort, but later arid population cohorts developed quicker than controls, indicating hormesis, an effect still present in the great-great-grandoffspring. These results demonstrate the transgenerational inheritance of a potentially beneficial hormetic phenotype and associated patterns of gene expression in a natural insect population.

**Significance Statement:** Global warming will lead to heat stress for many organisms. Interactions between the epigenome (the machinery controlling gene expression) and transposable elements, (TEs; mobile elements within the genome) are likely to influence the response to stress. In transgenerational laboratory heat shock experiments we investigated how the epigenome and TEs affected gene expression and phenotypes of arid and temperate populations of *Drosophila melanogaster*. The arid population showed clearer upregulation of epigenome and gene expression and transgenerational inheritance of gene expression following heat shock, and we identified TE insertions associated with stress-responsive genes. We also observed potentially beneficial transgenerational phenotypic responses in this population. Integrating molecular systems and measuring relevant phenotypes improves our understanding of acclimatory and evolutionary responses to environmental stress.

## Introduction

Temperatures on Earth are increasing due to anthropogenic climate change, presenting an evolutionary challenge for many populations (Burke et al. 2018; Catullo et al. 2019). Higher average temperatures and more extreme events such as heat waves will lead to increasingly frequent stressful events that can act as strong selective pressures (Grant et al. 2017). The fundamental mechanisms underlying heat stress response, including the expression of heat shock proteins (HSPs), are evolutionarily conserved. However, *hsp* and other stress-responding genes are remarkably diverse and show great variability in their regulation (Chen et al. 2018), reflecting adaptation to different environmental conditions. Rates of evolutionary change are generally assumed to be dependent on the amount of standing genetic variation and mutation rate in the population (Orr 2005; Bomblies and Peichel 2022). Yet evidence is emerging that some organisms adapt to environmental change more rapidly than might be predicted by these factors alone (Colautti and Barrett 2013; Campbell-Staton et al. 2017; Crotti et al. 2021). Two key mechanisms that could facilitate rapid adaptation and deserve further investigation are environmental sensitivity in the epigenome and transposable element (TE) activity (Rey et al. 2016; Pimpinelli and Piacentini 2020; McGuigan et al. 2021).

The epigenome is a set of interacting chemical marks and molecules, including DNA methylation, histone modification and chromatin accessibility that maintain the genomic DNA’s structure (Bannister and Kouzarides 2011; Dabin et al. 2016) and regulate gene expression (Taudt et al. 2016; Adrian-Kalchhauser et al. 2020). This regulation can be associated with pre-programmed developmental stages or environmentally sensitive changes that maintain organismal function or promote context-dependent alternative physiological or developmental responses (Adrian-Kalchhauser et al. 2020). Many organisms also rely on epigenomic mechanisms to suppress and regulate the activity of TEs (Slotkin and Martienssen 2007; Hisanaga et al. 2023). TEs are selfish genetic elements that insert new copies of themselves into the genome, influencing the evolution of genome structure and gene regulation (Feschotte 2008; Hayward and Gilbert 2022). Novel TE insertions can influence gene expression not only when they insert into or near to genes, but also when epigenomic silencing of TEs spreads to surrounding areas of the genome (Huang et al. 2022; Coronado-Zamora and González 2023). Furthermore, TE activity appears to increase in stressful conditions (Casacuberta and González 2013; Fanti et al. 2017; Horváth et al. 2017), potentially as a consequence of competing demands for the host’s epigenomic machinery (Cappucci et al. 2019). This can lead to higher frequencies of insertions associated with stress-response genes (Rech et al. 2019).

As well as affecting how individuals respond to heat stress, both the epigenome and TEs have the potential to influence responses to heat stress over multiple generations. There is growing evidence that environmentally-induced epigenetic changes can be passed on to later generations in a phenomenon known as transgenerational epigenetic inheritance or TEI (Hu and Barrett 2017; Harney et al. 2022), whereby certain epigenetic marks, instead of being reset between generations (Kawashima and Berger 2014) persist across them (Ou et al. 2012), and shape phenotypic responses of descendants (Crotti et al. 2021). If parental and grandparental conditions are predictive of the current environment, TEI could facilitate local adaptation over short evolutionary timescales (Bonduriansky et al. 2012; Fitz-James and Cavalli 2022). Furthermore, novel TE insertions that arise due to heat stress could be a potent source of genetic variation in populations that regularly experience this stress (Fanti et al. 2017; Rech et al. 2019). Previously, evidence for epigenetically inherited transgenerational effects often came from laboratory strains with specific mutations or knock-downs (Wan et al. 2021), and experiments considering phenotypes that do not relate clearly to organismal fitness (Ciabrelli et al. 2017; Fanti et al. 2017). Thus we are only now beginning to appreciate the extent to which epigenetic responses to stress vary in natural animal populations (Baduel et al. 2024). Determining the relevance of these effects on rapid adaptation and evolution under global environmental change requires further studies that considers both naturally occurring genetic differences and phenotypically relevant traits.

To understand how population genomic variation in TEs and variation in the epigenome combine under heat stress and potentially generate transgenerational effects requires an integrative experimental approach that considers the effect of natural variants and relevant phenotypes (Catullo et al. 2019). Here we study the effect of acute heat shock on female *Drosophila melanogaster* from two ecologically distinct European populations. We characterise the effects of heat shock on chromatin accessibility and gene expression in the ovaries, where epigenetic effects are likely to play a pivotal role in preparing embryos for early development and also in preventing the incorporation of novel TE copies into the germline. We relate expression and accessibility differences to polymorphic TEs that differ between the two populations and investigate the phenotypic consequences of the heat shock on offspring development traits. We then explore the potential for transgenerational effects in these populations by investigating whether ancestral heat shock continues to affect gene expression, chromatin accessibility and phenotypic development after three generations. This integrative approach allows us to measure transgenerational effects of heat shock on molecular and organismal phenotypes, and to determine whether variation in chromatin accessibility and polymorphic TE insertions play any role in their transmission across generations.

## Results

### Heat shock effects on expression and accessibility differed between populations with different thermal tolerances

To determine whether flies from temperate (Akaa, Finland) and arid (Manzanares: Manz, Spain) climates differed in their thermal tolerance, we measured their critical thermal maximum (CT_Max_) in the F2. Flies from Manz had a higher CT_Max_ (Chisq = 8.87, df = 1, P = 0.0029; **Fig. 1A**), remaining active up to temperatures of 40.3°C, compared with 39.8°C in flies from Akaa. In the F3 generation we measured gene expression and chromatin accessibility changes in female ovaries from heat shock (HS) versus control (Ctrl) treatments for both populations, and looked for shared and unique differentially expressed genes (DEGs), and shared and unique genes associated with differentially accessible promoter regions (DARs). We found 2,287 DEGs shared by both populations (**Fig. 1B**, **Fig. S2A** and **S2B**), but heat shock induced different strengths of “unique” response between populations, with a much larger effect detected in Akaa (2,478 DEGs) than Manz (255 DEGs). Genes associated with DARs also showed some overlap, with 503 shared between populations (**Fig. 1C**; **Fig. S3A** and **S3B**). In contrast to the expression results, heat shock induced stronger unique effects on accessibility in Manz (1,575 DARs) than Akaa (625 DARs).

**Fig. 1.**
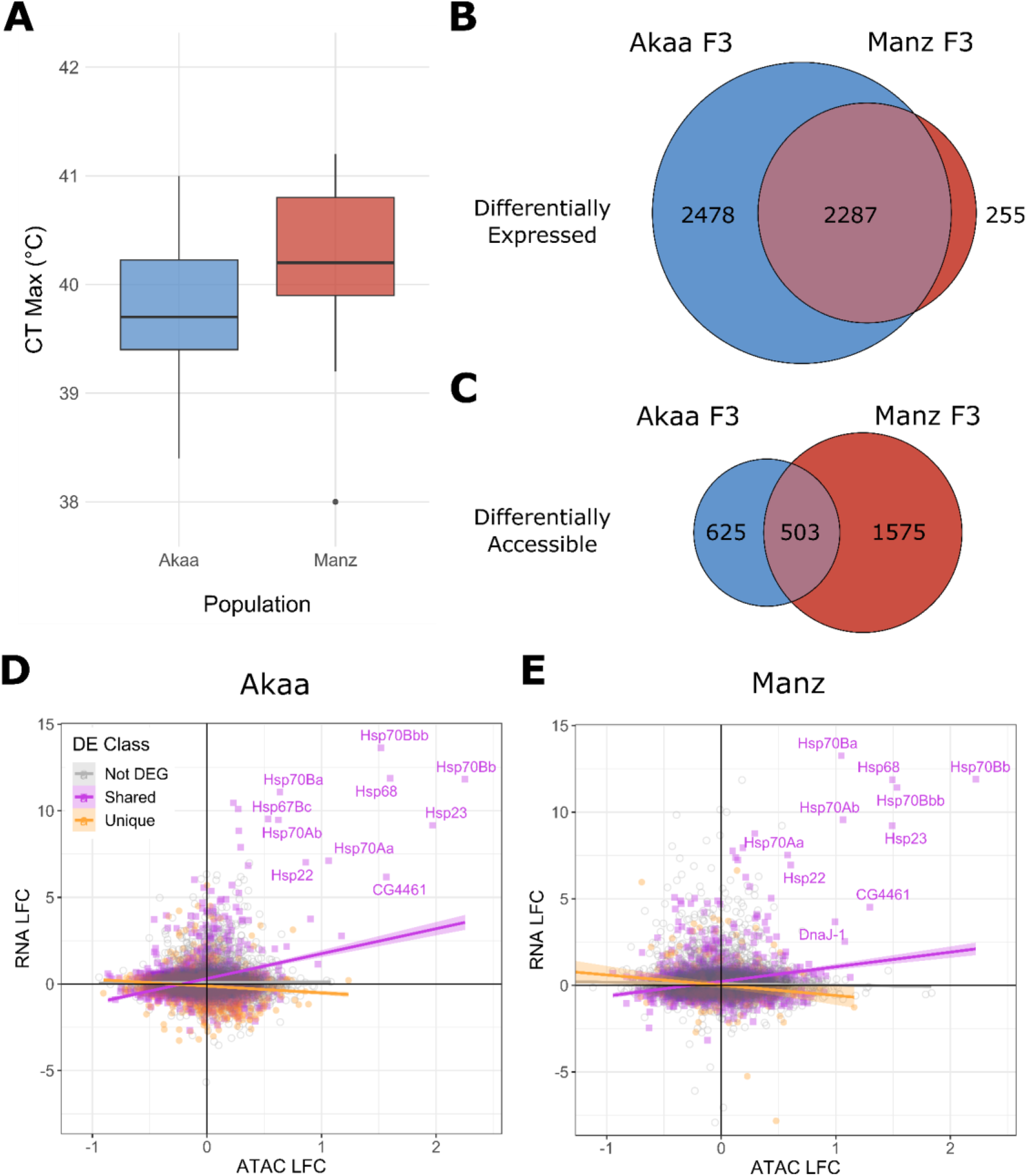
**A)** CT_Max_ of flies from Akaa and Manzanares. **B)** Overlap in differentially expressed genes (DEGs) in flies from Akaa and Manz, and **C)** Overlap in differentially accessible regions (DARs) in flies from Akaa and Manz. Correlation between log fold change (LFC) in RNA and LFC in ATAC following heat shock in **D)** Akaa and **E)** Manz. DEGs that overlap between populations are shared, those that don’t are unique. In **E**), two genes (CG32939 and P24-2) with strong negative ATAC and weak positive RNA values have been cropped to improve readability. The top 10 most correlated DEGs have been labelled: all are associated with heat shock proteins (HSPs), including CG4461 (part of the small HSP gene group), and DnaJ-1 (part of the Hsp40 gene group).

Because increased chromatin accessibility can promote increased gene expression, we expect some changes in these two measures to be concordant, indicating regulated gene expression. To determine the concordance between expression and accessibility changes following heat shock, we analysed RNA log fold-change (LFC) as a function of ATAC LFC, including DEG class (shared, unique, not DEG) as a main effect and through its interaction with ATAC LFC. RNA LFC was dependent on ATAC LFC in both Akaa and Manz, but the effect was highly dependent on the DE class (Akaa interaction: F = 220.41, df = 2, P < 0.0001; Manz interaction: F = 103.37, df = 2, P < 0.0001). In both Akaa (**Fig. 1D**) and Manz (**Fig. 1E**), there was a positive correlation between ATAC and RNA for shared DEGs, while unique DEGs showed negative correlations and non DEGs showed no (Akaa), or weak correlations (Manz). In both populations, concordance of expression and accessibility in shared DEGs was driven by large increases in expression and accessibility of heat shock proteins.

To further investigate the functional consequences of heat shock on gene regulation, we selected genes that showed both differential expression and differential accessibility of promoters (DE/DA genes). We found 447 DE/DA genes in Akaa and 426 in Manz. Among these genes, 114 were shared by both populations, 335 unique to Akaa, and 312 unique to Manz. Every single shared DE/DA gene showed the same pattern of regulation, i.e. those genes that showed concordant up-regulation in Akaa HS also showed concordant up-regulation in Manz HS. Two thirds of these shared DE/DA genes showed concordant expression and accessibility changes (**table 1**). For both populations less than half of the unique DE/DA genes showed concordant expression and accessibility changes, suggesting weaker gene regulation, but among genes that did show concordant expression and accessibility, a greater percentage showed up-regulation in Manz (19.23%) than Akaa (9.85).

**Table 1.** Percentages of genes that showed concordant and discordant patterns of expression and accessibility for shared and unique DE/DA genes in Akaa and Manz. All shared genes showed identical patterns of regulation in both populations. Exp: expression; Acc: accessibility.

|  | Shared | Akaa unique | Manz unique |
| --- | --- | --- | --- |
| Concord: Exp Up + Acc Up | 27.19 | 9.85 | 19.23 |
| Concord: Exp Dn + Acc Dn | 39.47 | 38.21 | 29.81 |
| Discord: Exp Up + Acc Dn | 17.54 | 37.31 | 25.64 |
| Discord: Exp Dn + Acc Up | 15.79 | 14.63 | 25.32 |

We then carried out functional enrichment analysis for different groups of DE/DA genes split by pattern of regulation (concordant up regulation, concordant down regulation, discordant regulation with expression up and accessibility down, and discordant regulation with expression down and accessibility up; **table 2**). Although we did not observe functional enrichment in all groups, shared DE/DA genes with concordant increases in expression and accessibility were highly enriched for functions relating to heat stress. More biological processes were functional enriched in uniquely DE/DA genes in Akaa (**table 2**), especially when accessibility was reduced (whether in combination with reduced expression or increased expression), which may suggest greater disruption of function following heat shock for the temperate population.

**Table 2.**
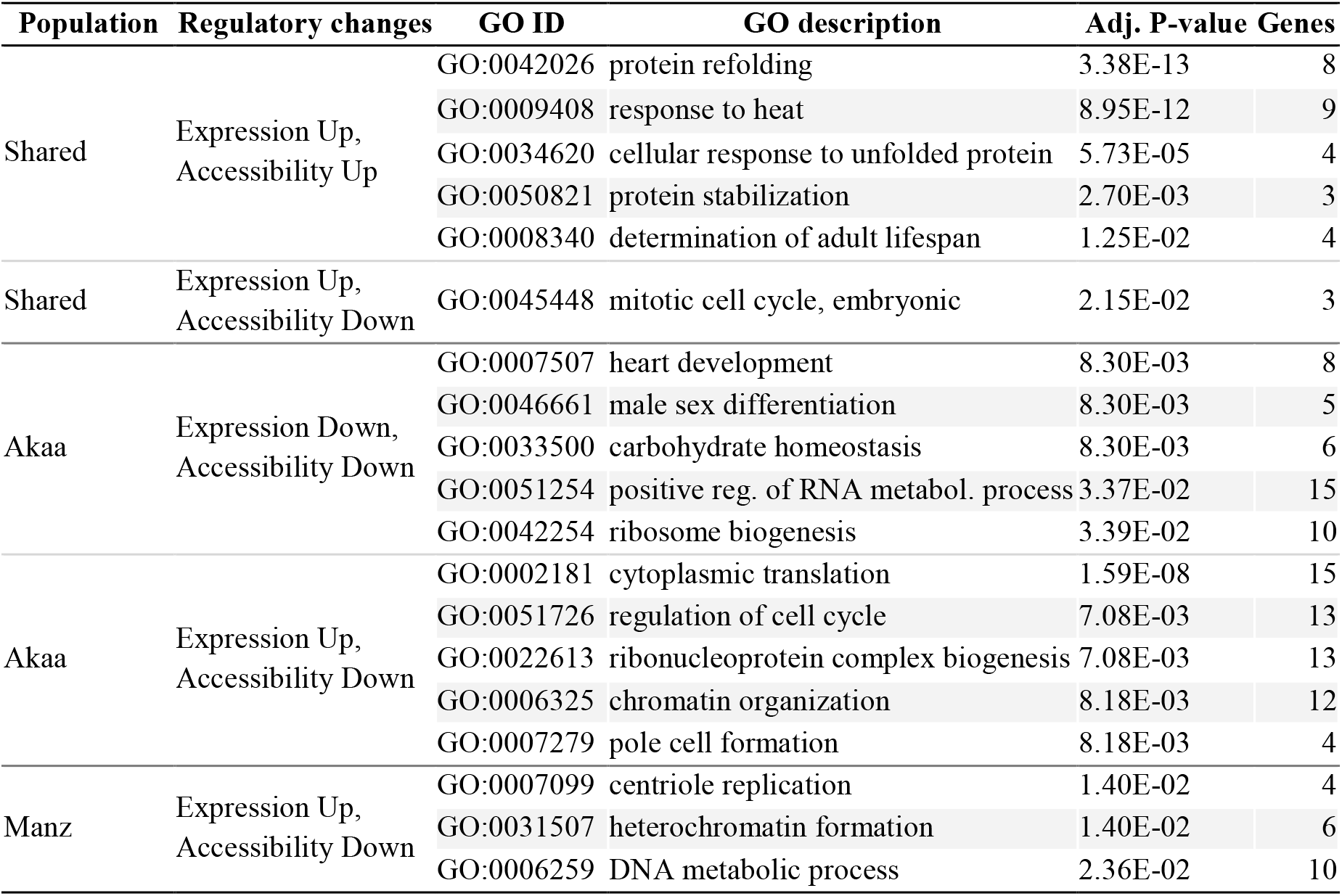
Functional enriched biological processes of genes that were differentially expressed and differentially accessible grouped by whether they were present in both populations or only one, and according to the concordance of expression and accessibility changes. Up to the top five biological processes are listed.

### Heat shock effects were transmitted across multiple generations, especially in the arid population

Heat shock continued to influence gene expression in both populations after three generations, with the effect stronger in Manz (292 DEGs) than Akaa (25 DEGs) (**Fig. 2A** and **2B**; **Fig. S2C** and **S2D**). Comparing the lists of heat shock DEGs in the F3 and the transgenerational DEGs in the F6 revealed minimal overlap between generations in Akaa (5 out of 25 F6 DEGs; **Fig. 2A**), but a larger overlap in Manzanares (132 out of 292 DEGs; **Fig. 2B**). Only four F6 DEGs (**Fig. 2C**) overlapped between populations (*MtnA*, *CG9953*, *Dph1*, and *CG18853*), none of which were F3 DEGs in either population.

**Fig. 2.**
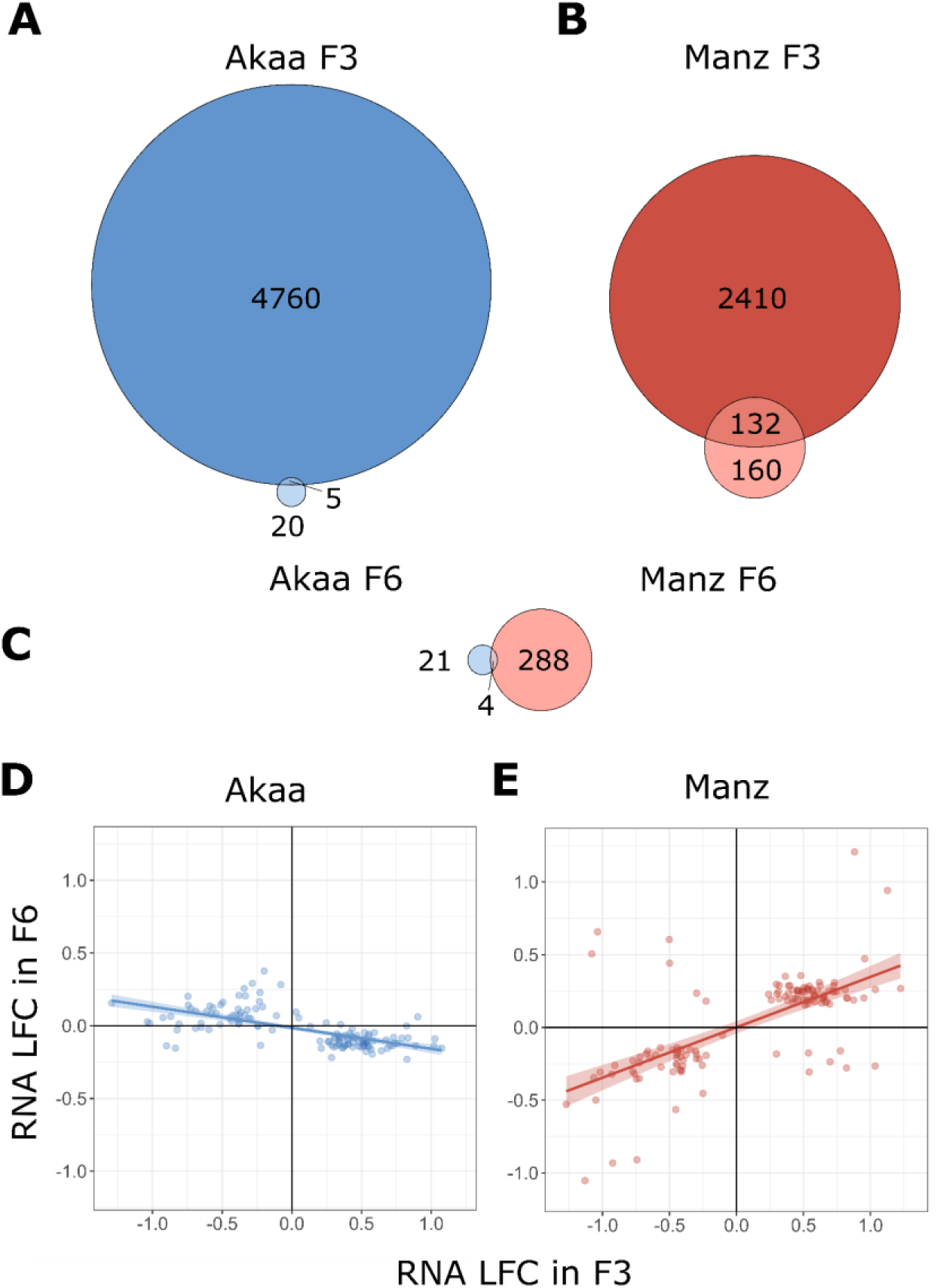
Overlap between DEGs directly responding to heat shock in the F3 and those that showed a response to ancestral heat shock in the F6 in **A)** Akaa and **B)** Manz. The overlap in F6 DEGs between the two populations is shown in **C)**. The correlation between F3 log fold change (LFC) and F6 LFC values are shown for a subset of DEGs in **D)** Akaa and **E)** Manz. This subset of genes were DEG in both F3 and F6 of Manz (i.e. the overlap in **B**), and 124 of them were DEG in the F3 of Akaa.

To see whether the 132 DEGs present in both Manz F3 and Manz F6 (**Fig. 2B**) were consistent in the direction of expression change between generations, we looked at the correlation in F3 and F6 LFC values. We could not repeat this approach for Akaa (due to the limited overlap in DEGs between generations); however, to provide a population comparison, we took the 132 transgenerational DEGs from Manz and looked at their F3 and F6 LFC values in Akaa: the majority of these genes (124/132) were differentially expressed in the F3 of Akaa (i.e. they were involved in the heat shock response), but none were differentially expressed in the F6. Gene expression was negatively correlated across generations in Akaa (coef. = −0.14, *F* = 97.96, *df* = 1, *P* < 0.0001; **Fig 2D**) and strongly positively correlated in Manz (*coef.* = 0.34, *F* = 98.24, *df* = 1, *P* < 0.0001; **Fig 2E**), with consistent expression direction changes in both generations for 90% (119/132) of genes in this population.

GO enrichment for the 132 DEGs present in both Manz F3 and Manz F6 (split into four groups based on the direction of expression in the 2 generations) found genes with consistent increases in expression following HS to be significantly enriched for biological processes including ‘mitotic cell cycle’, ‘nuclear division’ and ‘chromosome segregation’ (**table 3**), while genes with increased expression in F3 HS and decreased expression in F6 dHS may have had pentose or chitin metabolism functions.

**Table 3.** Functionally enriched GO biological process clusters for DEGs present in both Manz F3 and Manz F6. Terms are grouped by the direction of change in both generations (up = increase under heart shock, Dn = decrease under heat shock). For F3 Up, F6 Up, the top 10 GO processes are displayed.

| Transgenerational effect | GO ID | GO description | P-value | Genes |
| --- | --- | --- | --- | --- |
| F3 Up, F6 Up | GO:0000278 | mitotic cell cycle | 3.00E-16 | 27 |
|  | GO:0000280 | nuclear division | 1.03E-12 | 20 |
|  | GO:0007059 | chromosome segregation | 3.33E-07 | 12 |
|  | GO:0051276 | chromosome organization | 3.57E-07 | 14 |
|  | GO:0071824 | protein-DNA complex subunit organization | 2.10E-05 | 8 |
|  | GO:0051301 | cell division | 2.97E-05 | 11 |
|  | GO:0006260 | DNA replication | 2.97E-05 | 8 |
|  | GO:0006325 | chromatin organization | 5.30E-05 | 11 |
|  | GO:0007051 | spindle organization | 5.52E-05 | 8 |
|  | GO:0035220 | wing disc development | 6.53E-03 | 9 |
| F3 Down, F6 Up | GO:0019321 | pentose metabolic process | 3.87E-02 | 1 |
|  | GO:0006032 | chitin catabolic process | 3.87E-02 | 1 |
|  | GO:0018990 | ecdysis, chitin-based cuticle | 3.87E-02 | 1 |
|  | GO:0003014 | renal system process | 3.98E-02 | 1 |

We also inspected functional information from Flybase (https://flybase.org/) relating to DEGs induced by ancestral heat shock to identify genes associated with stress response or epigenome functions (searching for the key words ‘stress’; and ‘epigen’, ‘chromatin’, ‘histone’, and ‘methyl’). Among the four genes that were differentially expressed in both populations were a stress response gene (*MtnA*) and a putative tRNA methyltransferase (*CG18853*). In Akaa we found only two genes with potential stress response functions (**table S1)**, neither of which were differentially expressed in the F3. On the other hand, in Manz, we found 13 genes relating to stress response, 45 to the epigenome, and two with a potential function in both (*Hsf* and *Bicra*). Many of these genes (three related to stress response, 22 to the epigenome, and *Hsf*) showed transgenerational inherited patterns of expression, i.e. they were differentially expressed in both F3 and F6, with the direction of expression change consistent between generations (**table S2)**.

In contrast to the RNA-seq results, we found few transgenerational effects of heat shock on chromatin accessibility (**Fig. S3C** and **S3D**). Zero DARs associated with ancestral heat shock were observed in Akaa F6, and just three were found in Manzanares: *CG7966* (predicted to have methanethiol oxidase and selenium-binding activity), *pic* and *tara*. Although *tara* differed between generations in controls, it was also the only gene that was differentially accessible in both F3 and F6 (less accessible in F3 HS and more accessible in F6 dHS), and so merits consideration. *Tara* is thought to mediate the functions of trithorax group (TrxG) and polycomb group (PcG) genes (Dutta and Li 2017), potentially regulating transcription during development. To see if other TrxG or PcG genes showed transgenerational expression responses, we checked differential expression results in the F6 for 44 genes classified as ‘Trithorax group’ and 19 genes classified as ‘Polycomb group’ on Flybase (**table S2**). We found five TrxG genes (but no PcG genes) among F6 DEGs in Manz: *brm*, *mor*, *Bap111*, *Bicra*, and *nej* all showed increased expression in F6 dHS, and four of them (*brm*, *mor*, *Bap111*, and *nej*) showed transgenerationally inherited patterns of expression (direction of expression change consistent between generations). No TrxG or PcG genes were differentially expressed in the F6 of Akaa.

### TEs were associated with reduced expression in the temperate population and increased accessibility in the arid population

Next we considered whether F3 DEGs and DARs showed any associations with transposable elements (TEs). In Akaa we identified 1,559 reference insertions and 677 non-reference insertions, of which, respectively, 1087 and 440 were within 1kb of annotated genes. In Manz 1,659 reference and 609 non-reference insertions were identified, of which 1158 and 424 were within 1kb of annotated genes (**tables S3 and S4**). The presence of non-reference TEs is of particular interest, as these are more likely to be recent polymorphic insertions that might contribute to population differences.

Associations between TEs and patterns of expression and accessibility in the F3 were evaluated with Chi-squared tests. F3 DE class (not DE, shared, unique) was negatively associated with non-reference TEs in both populations (Akaa: Chisq = 16.66, P = 0.0023; Manz: Chisq = 19.20, P = 0.0007). On the other hand, F3 DAR class was positively associated with reference TEs in Akaa (Chisq = 16.07, P = 0.0029), but there was no association in Manz (Chisq = 6.06, P = 0.1949). In terms of the direction of change, the presence of non-reference TEs was positively associated with reduced expression in Akaa (Chisq = 18.07, P = 0.0001; **Fig. 3A**) but there was no association in Manz (Chisq = 2.05, P = 0.3588; **Fig. 3B**). On the other hand, TE presence was not associated with chromatin accessibility direction in Akaa (Chisq = 1.78, P = 0.4098; **Fig. 3C**), but both reference and non-reference TEs were associated with increased accessibility in Manz (Chisq = 38.54, P < 0.0001; **Fig. 3D**).

**Fig 3.**
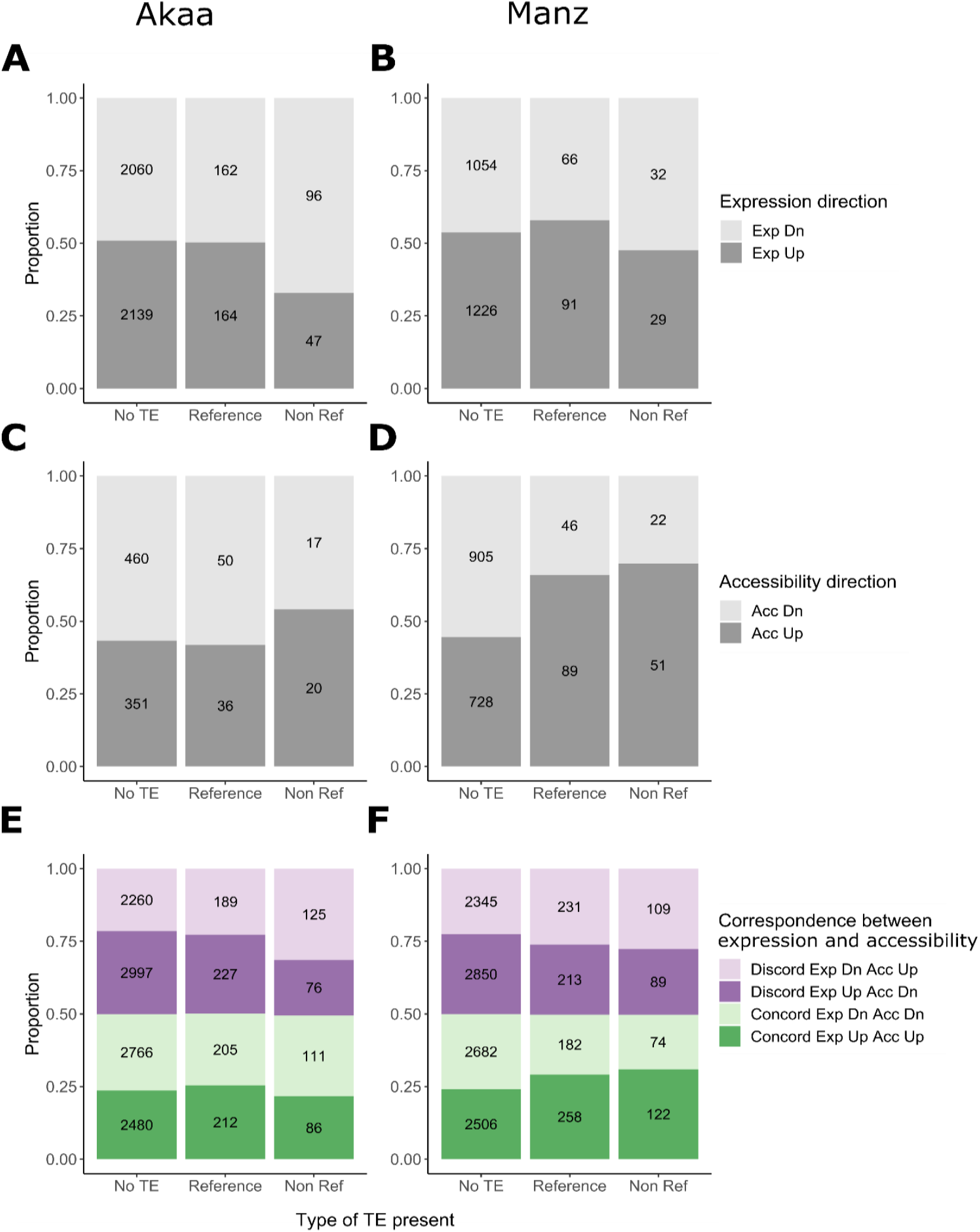
Associations between TE presence and genes with significant expression changes in the F3 in **A)** Akaa and **B)** Manz, and associations between TE presence and genes with significant accessibility changes in the F3 in **C)** Akaa and **D)** Manz. Effect of TE presence on the proportion of concordantly and discordantly regulated genes in response to F3 heat stress in **E)** Akaa and **F)** Manz.

Considering more general associations between gene regulation and TE presence using concordance in expression and accessibility changes for all genes (not just DEGs and DARs), we found associations in both populations. In Akaa, non-reference TEs were positively associated with genes that had increased accessibility but reduced expression (Chi.sq = 32.85, P < 0.0001; **Fig. 3E**). In Manz all TEs were positively associated with genes with increased expression and accessibility, and negatively associated with reduced expression and accessibility (Chi.sq = 48.05, P < 0.0001; **Fig. 3F**). Results of post-hoc tests for all significant Chi-squared tests are in **table S5**.

To further investigate the links between TEs and differential expression (Akaa) or accessibility (Manz), we carried out functional enrichment analysis on groups of differentially expressed or accessible genes proximate TEs. For Akaa, DEGs with reduced expression near to non-reference TEs were enriched for biological processes including ‘cellular response to endogenous stimulus’ (GO:0071495, adj. P = 0.0005, 10 genes) and ‘neuron projection development’ (GO:0031175, adj. P = 0.0005, 14 genes). In Manz, DARs with increased accessibility near to non-reference TEs were enriched for ‘cell-cell adhesion’ (GO:0098609, adj. P = 0.005, 6 genes) and ‘regulation of receptor-mediated endocytosis’ (GO:0048259, adj. P = 0.025, 3 genes), while those near to reference TEs were enriched for ‘ommatidial rotation’ (GO:0016318, adj. P = 0.01, 4 genes) and ‘sensory organ development’ (GO:0007423, P. adj = 0.01, 12 genes). Full lists of enriched GO terms can be found in **table S6**.

To identify candidate TEs linked to population differences in gene expression and chromatin accessibility following heat shock, we compiled lists of polymorphic TEs that were present only in one population, and near to genes that were DE or DA in that population. In Akaa we found 44 genes associated with TEs unique to that population: 36 with differential expression, four with differential accessibility, and four with both expression and accessibility changes (**table S7**). The top 10 DEGs by absolute LFC are shown in **table 4**. In Manz we found 20 genes associated with TEs unique to that population: three with differential expression, 16 with differential accessibility and one with both expression and accessibility changes (**table S8**). The top 10 DARs by absolute LFC are shown in **table 5**. We also found one gene in Akaa and 11 in Manz that were associated with unique TEs and unique differential expression in the F6 generation (**table 6**).

**Table 4.** Top 10 genes for which a TE insertion was proximate (within 1 kb) in Akaa but not Manz, and which were differentially expressed in Akaa F3 but not Manz F3 following heat shock (HS). Genes are ordered by absolute LFC value (Npc2g and Npc2h are grouped together due to being proximate to the same TE insertion). Significant LFC values are in bold.

| Transposable Element Insertion |  |  | Gene Associated with TE |  |  |  |  | Akaa F3 HS vs Ctrl |  | Manz F3 HS vs Ctrl |  |
| --- | --- | --- | --- | --- | --- | --- | --- | --- | --- | --- | --- |
| TE family | Chr. | Insertion Range | Type* | Symbol | Strand | TE Position | Response | RNA LFC | ATAC LFC | RNA LFC | ATAC LFC |
| 412 | 3R | 30744578 - 30744582 | NR | *Npc2g* | -1 | downstream | Both | **-1.69** | **0.466** | -0.46 | -0.079 |
|  |  |  |  | *Npc2h* | -1 | downstream | Exp | **-1.396** | 0.269 | -0.451 | -0.213 |
| I-element | 2R | 12927664 - 12927809 | NR | *sug* | 1 | intronic | Exp | **-1.582** | -0.023 | -0.604 | -0.268 |
| Jockey | 3L | 7130748 - 7130903 | NR | *Acbp2* | 1 | upstream | Exp | **-1.363** | 0.185 | -0.398 | 0.194 |
| BS | 2R | 18807728 - 18807877 | NR | *CG15096* † | -1 | downstream | Exp | **-1.116** | -0.01 | -0.602 | 0.017 |
| Roo | 3R | 13653876 - 13653953 | NR | *rin* | 1 | intronic | Exp | **-0.664** | -0.236 | -0.436 | -0.268 |
| I-element | 3R | 14520763 - 14520812 | NR | *Pde6* | -1 | intronic | Exp | **-0.58** | 0.251 | -0.349 | 0.12 |
| Jockey | X | 11476104 - 11476207 | NR | *Cyp4g15* | 1 | upstream | Both | **0.467** | -0.556 | 0.439 | -0.223 |
| Jockey | 3R | 11828464 - 11828542 | NR | *CG6959* † | -1 | intronic | Exp | **-0.435** | 0.1 | -0.345 | 0.268 |
| Jockey | 2L | 12382257 - 12382336 | NR | *bru2* | 1 | intronic | Exp | **-0.432** | 0.068 | -0.252 | 0.247 |

\* Type: was the insertion a reference (R) or non-reference (NR) insertion.
† *CG15096* is predicted to belong to the *SLC17* family of organic anion transporters; *CG6959* is orthologous to human *TPBG* (trophoblast glycoprotein).

**Table 5.**
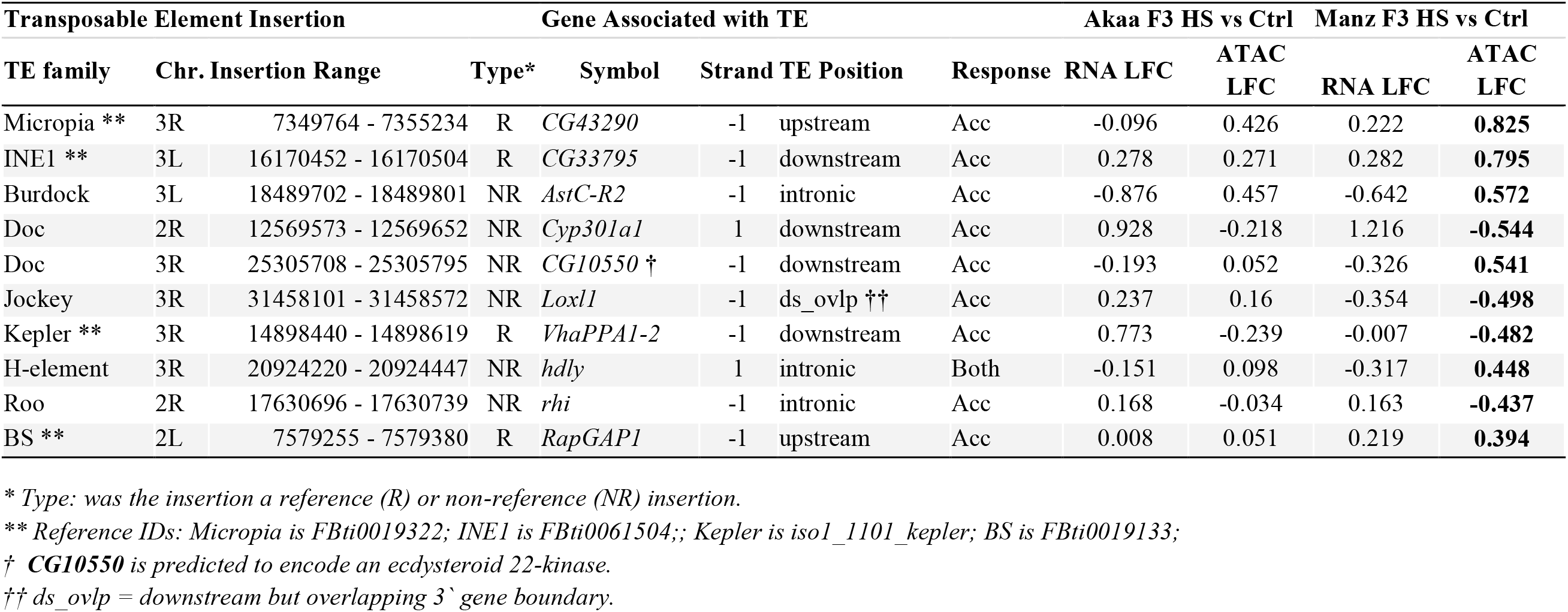
Top 10 genes for which a TE insertion was proximate (within 1 kb) in Manz but not Akaa, and which were differentially accessible in Manz F3 but not Akaa F3 following heat shock (HS). Genes are ordered by absolute LFC value. Significant LFC values are in bold.

**Table 6.**
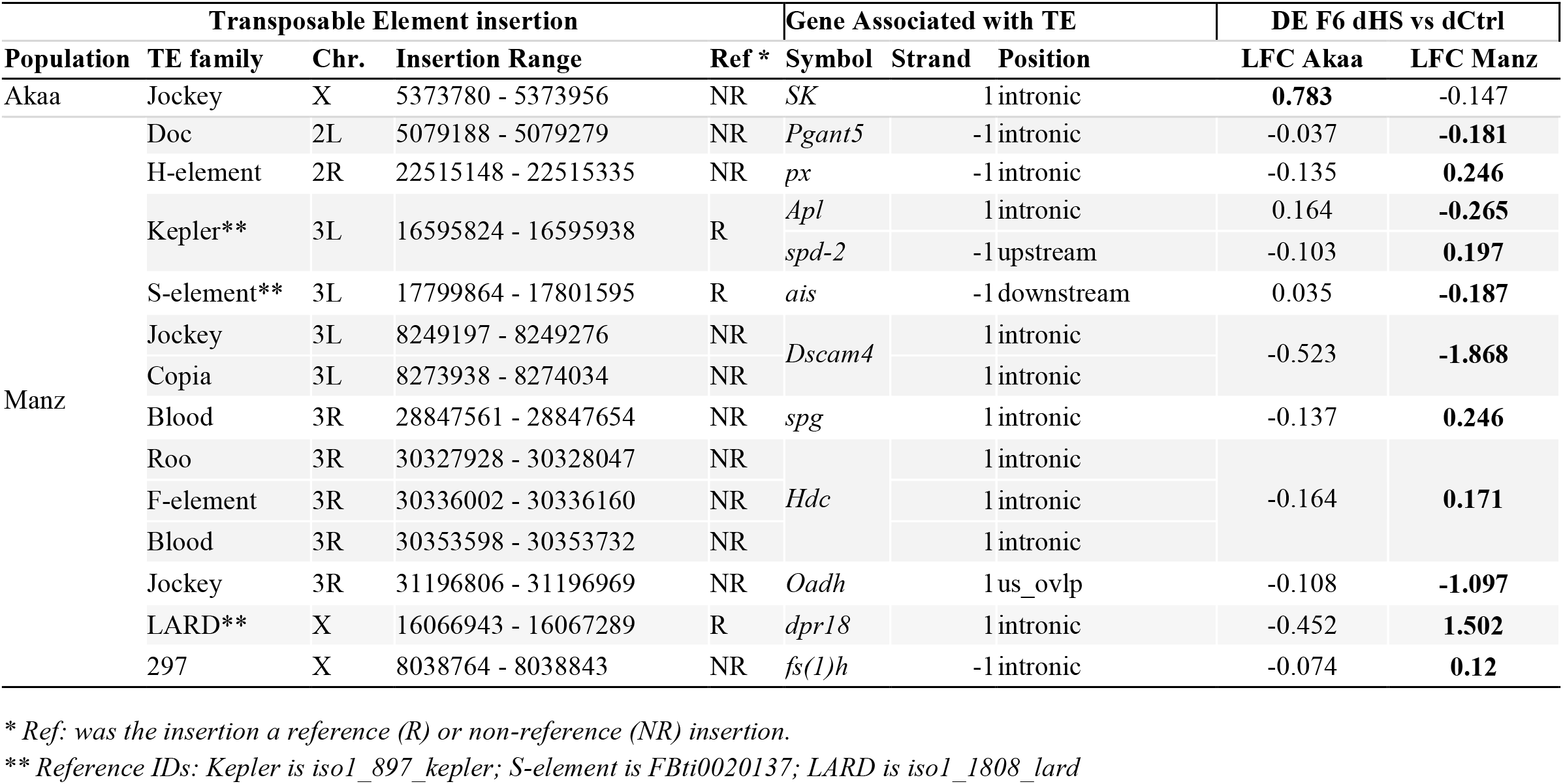
Polymorphic TE insertions and their associated genes that were differentially expressed in F6 dHS in one population and not the other.

Finally, we looked at the list of genes that were differentially expressed transgenerationally (DEG in both F3 and F6) to see how many were associated with TE insertions. For Akaa a single transgenerationally expressed gene (*CG43333*) was associated with a reference TE insertion, but this insertion was present in both populations. In Manzanares, we found 13 transgenerational DEGs associated with TEs. For 11 of these, the insertions were not polymorphic, but for two genes (*px* and *spd-2*), the insertions appeared to be unique to Manz (**table S9**). A single table combining expression, accessibility and TE results across generations for both populations is provided in **table S10**.

### Heat shock induced direct phenotypic effects and transgenerational effects in the arid population

We quantified the phenotypic consequences of F3 heat shock for the F4 offspring and F7 great-great-grand-offspring by measuring the number of eggs, pupae, and adults, the egg-to-pupa and egg-to-adult viability, and the times to pupation and eclosion. For the first cohort of F4 (eggs laid within two days of treatment), heat shock resulted in reduced numbers of eggs (*Chisq* = 13.11, *df* = 1, *P* = 0.0003), pupae (*Chisq* = 116.02 *df* = 1, *P* < 0.0001) and adults (*Chisq* = 165.59, *df* = 1, *P* < 0.0001) in both populations (**Fig. S4**). It also led to reduced egg-to-pupa viability (*Chisq* = 72.04, *df* = 1, *P* < 0.0001; **Fig. S5A**) and egg-to-adult viability (*Chisq* = 74.57, *df* = 1, *P* < 0.0001; **Fig. 4A**), and population-dependent delays in time to pupation (Trt x Pop: *F* = 11.830, *df* = 1, *P* = 0.0006; **Fig. S5B**) and time to eclosion (Trt x Pop: *F* = 11.02, *df* = 1, *P* = 0.0009; **Fig. 4B**). Post-hoc tests revealed the effects on time to pupation and eclosion to be stronger in Akaa (pupation: *t* = -7.047 *P* <0.0001; eclosion *t* =−6.864, *P* < 0.0001) than Manzanares (pupation: *t* =−2.894, *P* = 0.0039; eclosion *t* =−2.857 *P* = 0.0044).

**Fig 4.**
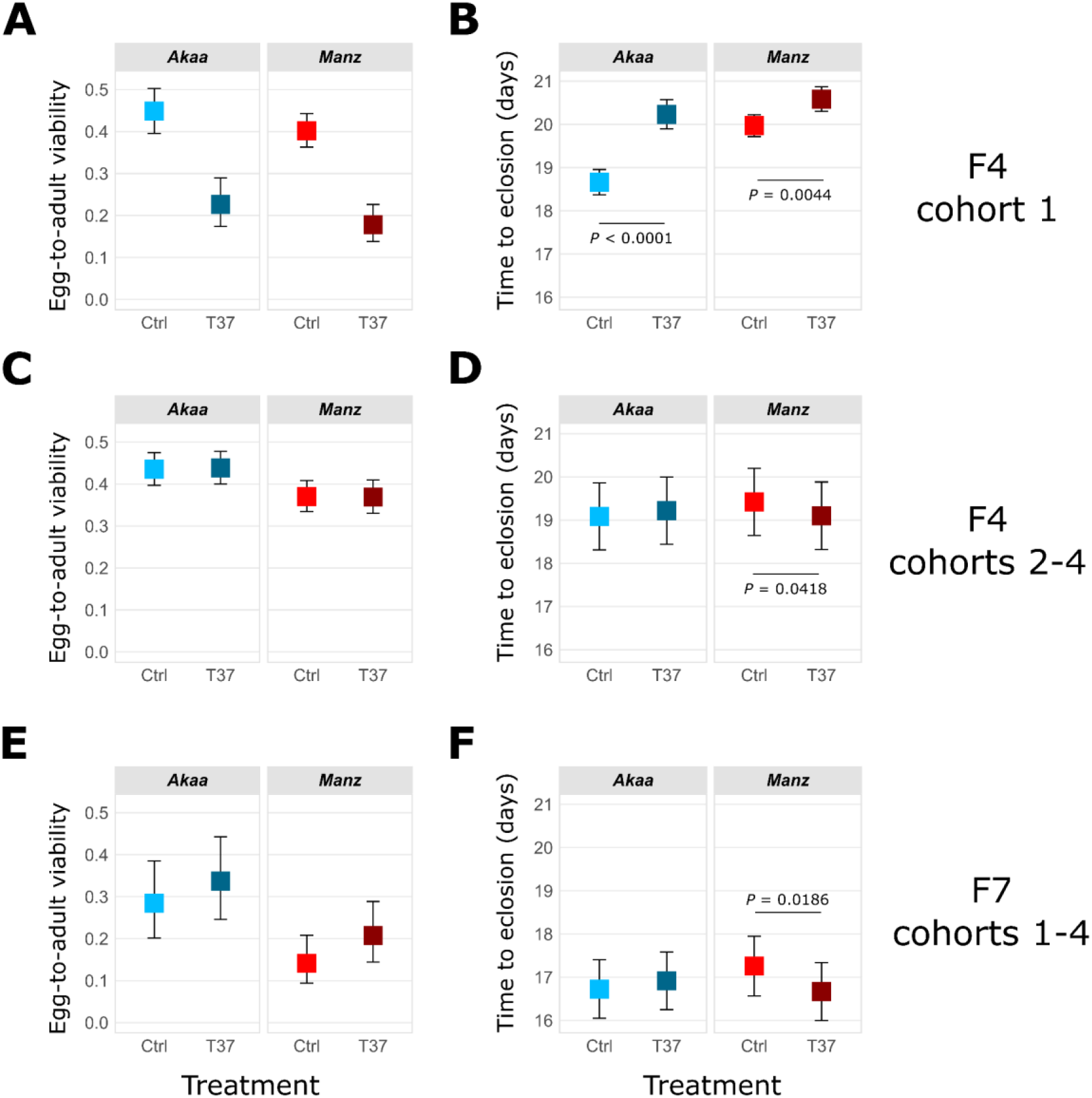
Effects of F3 heat shock on offspring and great-grand-offspring phenotypes. **A)** Egg-to-adult viability, and **B)** time to eclosion in F4 offspring from the first cohort, produced within 48 hours of treatment of F3 females (cohort 1). **C)** Egg-to-adult viability, and **D)** time to eclosion in F4 offspring from cohorts 2-4, produced in 3 separate 48 hour periods between 2 and 14 days after treatment (cohorts 2-4). **E)** Egg-to-adult viability, and **F)** time to eclosion in the F7 great-grand-offspring of F3 heat shocked females from all four cohorts. Results of post-hoc tests for population-specific treatment effects are shown within panels.

In later cohorts of F4, negative effects of heat shock on absolute numbers and viabilities were not significant (numbers of eggs, pupae, and adults, **Fig. S5**; egg-to-pupa viability, **Fig. S6A**; egg-to-adult viability, **Fig. 4C**), suggesting that females rapidly recovered from the heat shock. However there were population-specific effects of heat shock on time to pupation (Trt x Pop: *Chisq* = 6.0516, *df* = 1, *P* = 0.0139; **Fig. S6B**) and time to eclosion (Trt x Pop: *Chisq* = 4.5598, *df* = 1, *P* = 0.0327; **Fig. 4D**). In both cases we observed reduced development time for heat shocked flies from Manzanares (pupation: *t* = 3.087, *P* = 0.0027; eclosion: *t* = 2.065, *P* = 0.0418), indicating a hormetic effect.

Heat shock also resulted in transgenerational phenotypic effects in the F7. Although there was no effect of ancestral heat shock on the numbers of eggs, it did lead to a slight increase in the number of pupae (*Chisq* = 4.6056, *df* = 1, *P* = 0.0319) and adults (*Chisq* = 3.8631, *df* = 1, *P* = 0.0494) (**Fig. S7A-C**). This led to an increase (albeit statistically non-significant) in egg-to-pupa and egg-to-adult viability for both populations (egg-to-pupa: *Chisq* = 3.3119, *df* = 1, *P* = 0.0688, **Fig. S7D**; egg-to-adult: *Chisq* = 2.5663, *df* = 1, *P* = 0.1092, **Fig. 4E**). Furthermore, the population-specific reduction in time to eclosion that was observed in F4 cohorts 2-4 was recapitulated in the F7 (Trt x Pop: *Chisq* = 5.4545, *df* =1, *P* = 0.0195; **Fig. 4F**), i.e. descendants of heat shocked flies from Manz eclosed earlier (*t* = 2.428, *P* = 0.0180). The effect on time to pupation was in the same direction, but the interaction was not significant (Trt x Pop: *Chisq* = 2.4659, *df* = 1, *P* = 0.1163434; **Fig. S7E**).

## Discussion

Heat stress will increasingly affect many organisms as climate change leads to higher ambient temperatures and more extreme events such as heat waves. To understand how populations might acclimate and potentially adapt to these stressful environments, we explored interactions between transcriptional and epigenetic responses to heat shock, their associations with TEs, and their phenotypic consequences in two populations of *D. melanogaster*, and investigated whether any aspects of the heat shock response were transgenerationally inherited. We found many differences in how heat shock responses were regulated between populations, including important associations between TEs and expression and chromatin accessibility, and transgenerational phenotypes indicative of hormesis in the arid population.

### Concordant changes in expression and accessibility following heat stress were more coordinated in the arid population

Heat shock induced many changes in *D. melanogaster* ovary gene expression and chromatin accessibility in both populations. Focusing on genes that were both differentially expressed and differentially accessible (DE/DA) in both populations, we observed a robust and concordant increase in heat shock proteins (HSPs), which are key regulators of the stress response (Feder and Hofmann 1999; Saibil 2013), reflected in the functional enrichment of GO biological processes such as ‘response to heat’, and ‘protein refolding’. When considering population-specific responses, there were more concordant increases in DE/DA genes (and fewer concordant decreases) in the arid population than the temperate population, which could suggest a better coordinated response to the stress in the population that experiences heat shock more frequently (Vihervaara et al. 2018). However, in both populations just over half of the DE/DA genes showed discordant patterns of expression and accessibility, highlighting that while chromatin accessibility changes might facilitate transcriptional changes, they probably do not drive them (Ing-Simmons et al. 2021). Furthermore, discordance between these two molecular phenotypes may be a consequence of the environmental stress itself (Kiani et al. 2022).

### Transcription but not chromatin accessibility was transgenerationally inherited in the arid population

Heat shock continued to influence gene expression three generations after the treatment, with 292 DEGs observed in the descendants of heat shocked flies from the arid population, including 132 DEGs that responded in both F3 and F6 generations. These DEGs positively correlated across generations, indicating transgenerational epigenetic inheritance (TEI). Our understanding of the molecular factors influencing TEI in animals has grown in recent years (Ciabrelli et al. 2017; Kishimoto et al. 2017; Wan et al. 2021), and among the transgenerationally inherited DEGs in the arid population were critical genes identified from previous TEI experiments in *C. elegans*, such as *Hsf* (Kishimoto et al. 2017), and a SET domain histone methyltransferase (Woodhouse et al. 2018; Wan et al. 2021). In a recent genome-wide association study and functional validation of stress preconditioning in *D. melanogaster*, Owings and Chow (2024) found that Set1 was critical to establishing stress memory; furthermore, among their shortlist of candidate genes for stress preconditioning were four which showed TEI in this study (*CG7781*, *CG16812*, *px* and *Drat*).

Epigenetic genes are thought to help establish environmentally induced TEI (Frolows and Ashe 2021), and we identified 23 genes with likely epigenetic functions, including four belonging to the trithorax group (TrxG) of genes (*brm*, *Bap111*, *mor*, *nej*). TrxG genes activate transcription and interact with repressive PcG proteins to regulate the cell cycle and development (Schuettengruber et al. 2011), stress resistance (Siebold et al. 2010) and maintenance of epigenomic memory (Geisler and Paro 2015). Two of these TrxG genes (*Bap111*, *mor*), together with the chromatin remodelling genes *Chd1* and *Chd3* (also TEI in the arid population), are involved in SWI/SNF ATP-dependent chromatin remodelling, which helps regulate cell proliferation (Tian and Smith-Bolton 2021), and has been implicated in thermal tolerance (Ji et al. 2022). These genes thus represent interesting candidates for further research into TEI and hormetic responses to heat stress.

Although many epigenetic genes were differentially expressed, we actually observed few transgenerational effects of heat shock on chromatin accessibility itself: three genes were more accessible in the descendants of heat shocked flies in the arid population (*CG7966*, *pic*, and *tara*). The transcriptional co-regulator, *tara*, may be of interest as it mediates interactions between TrxG and PcG proteins during chromatin remodelling and cell fate determination (Calgaro et al. 2002; Schuster and Smith-Bolton 2015; Dutta and Li 2017). However, given the small number of transgenerational changes in chromatin accessibility we observed, it is possible that transgenerational changes in transcription were driven by other mechanisms such as germline transfer of small RNAs (Ashe et al. 2012; Wilson et al. 2023) or long-range chromatin interactions (Ciabrelli et al. 2017).

### TEs were associated with discordant regulation in the temperate population but concordant regulation in the arid population

A key factor that could underlie population differences in the regulation of the stress response is the presence of transposable elements (TEs), which have the ability to rewire regulatory networks (Chuong et al. 2017) and regulate stress-responsive genes (Horváth et al. 2017). In the temperate population (Akaa), non-reference TEs were associated with reduced expression but not reduced accessibility, suggesting that other mechanisms such as piRNA expression (Siomi et al. 2011) may suppress the expression of certain genes close to TEs. In the arid population (Manz) on the other hand, TEs were associated with genes with significantly more accessible chromatin but not increased expression, which could reflect preferential insertion into regions of accessible chromatin (Cao et al. 2023), and highlights that epigenetic changes linked to TEs may not always result in the expected transcriptional changes (Coronado-Zamora and González 2023). Our results contribute to growing evidence that the effects of TEs on the epigenome and gene expression vary among populations and contribute to gene regulatory evolution (Coronado-Zamora and González 2023; Bodelón et al. 2023).

Furthermore, TE-associated genes with reduced expression (Akaa) and increased accessibility (Manz) were functionally enriched for numerous developmental GO biological processes (**table S6**), highlighting the potential for TEs to influence gene-regulation during development (Todd et al. 2019). Looking more closely at lists of polymorphic (population-specific) TEs associated with unique DEGs or DARs also reveals interesting candidates. For example, in Akaa (**table 4**), the top three genes influenced by polymorphic TEs were *Npc2g*, *Npc2h* and *Sug*, which are all involved in the lysosome and autophagy, suggested as a mechanism of heat-stress tolerance in *D. melanogaster* (Willot et al. 2023). Similarly, in Manz (**table 5**) increased expression of specific candidates bears further investigation. *AstC-R2*, which regulates egg production following cold-induced reproductive dormancy (Meiselman et al. 2022) and *Rhi,* which is involved in the piRNA-mediated transposon repression pathway.

In Manz we also observed two genes with transgenerational expression that were proximate to polymorphic TE insertions (**table 6**). S*pd-2* plays a critical role in centrosome maturation during mitosis (Wong et al. 2024), while *px* is known to influence wing morphogenesis (Matakatsu et al. 1999), and as mentioned previously, was a candidate gene for stress preconditioning (Owings and Chow 2024). Other transgenerationally expressed genes in Manz which were proximate to (non-polymorphic) TEs may be worthy of further study: *Dscam4*, for example, is thought to affect thermosensation (Corthals et al. 2023). The aforementioned candidates could all provide useful starting points to investigate how TEs might shape gene regulatory responses to heat stress within and across generations.

### Flies recovered from heat shock and those from Manzanares showed a transgenerational hormetic response

Heat shock not only affected chromatin accessibility and gene expression in the F3, but had a strong immediate effect on offspring phenotype in the F4, and subtle effects on F4 offspring that were transgenerationally inherited by the great-grand-offspring in the F7. Strong negative effects on viability and development time were observed in the first cohort of offspring from both populations, as shown in classic heat shock experiments (Krebs and Loeschcke 1994; Silbermann and Tatar 2000). Yet negative effects all but disappeared in eggs laid more than two days after the heat shock, suggesting that females largely recovered from the heat shock in the medium-term (two to 14 days). The initial deleterious effects of heat shock could have been caused by damage to the females’ sperm reserves, which are susceptible to heat stress (Sales et al. 2018; Iossa 2019), or damage to partially-developed embryos retained within the reproductive tract (Horváth and Kalinka 2018).

Furthermore, heat shocked females from Manzanares produced offspring with more rapid development, indicating a possible hormetic effect. Hormesis occurs when a low dose of stress leads to improved physiological function later in life (Costantini et al. 2010). In *Drosophila*, hormesis can influence reproduction and survival (Le Bourg et al. 2001; Rix and Cutler 2022) and rates of development (Zhou et al. 2020). These effects can differ among genotypes (Gomez et al. 2016), and in line with our results, a recent study in *D. buzzatii* found that hormetic responses to heat stress were stronger in heat-tolerant populations (Almirón et al. 2024). More unusually, the hormetic effect of heat shock on development time was still present in Manz after three generations. Although beneficial hormetic maternal effects (Margus et al. 2019) and disadvantageous transgenerational phenotypic effects (Mu et al. 2021) have previously been observed in insects, to our knowledge this is the first observation of a potentially beneficial transgenerational hormetic phenotypic effect operating over more than two generations in a natural insect population.

## Conclusions

Our results show that heat shock induced strong changes in gene expression and chromatin accessibility, and that upregulatory responses were stronger in the arid population. Changes in expression in the temperate population, and accessibility in the arid population, were also associated with the presence of TEs. The arid population displayed transgenerational inheritance of gene expression in genes previously identified to facilitate *C. elegans* transgenerational effects such as *hsf* and SET-domain histone methyltransferase, as well as many genes with cell cycle process and epigenome modifying functions. These findings support the idea that chromatin remodelling may facilitate TEI (Ruden and Lu 2008; Sabarís et al. 2023), although we did not find strong evidence for heat shock influencing chromatin accessibility transgenerationally. TEI was accompanied by a potentially beneficial hormetic phenotypic response in the offspring and the great-great-grand-offspring of heat shocked flies from the arid population, and we hypothesise that heat shock induced changes in rates of cell proliferation that were transgenerationally inherited across at least three generations. Increased speed of larval and pupal development may allow flies to limit their time in necrotic fruit which can frequently overheat in natural conditions (Feder et al. 1997), and our results provide evidence that environmentally induced changes in the epigenome may target certain genes to generate transgenerational developmental plasticity. This effect could grant transgenerational hormesis an evolutionary role, linking acclimation with adaptation (Costantini, 2019).

## Materials and Methods

### Fly lines and experimental overview

Wild *D. melanogaster* (F0) were collected from one site in Spain near the town of Manzanares, abbreviated to Manz (38.98°N, 3.35°W), and another site near to Akaa in Finland (61.10°N, 23.52°E) in late September 2021. For each population, offspring from 10 F0 females were selected to set up F1 lab populations. We established large embryo collection cages with 100 (Manz) or 150 (Akaa) F1 females and a similar number of F1 males. The diversity of F0 females was equally represented in the number of F1 female founders, i.e. for Manz 10 F0 each contributed 10 F1 females and for Akaa 10 F0 each contributed 15 F1. Large lab populations were maintained in cages until the F3, but experimental animals (from F3 to F7) were maintained in standard *Drosophila* tubes. Flies were kept in the lab, with natural fluctuations in temperature and light:dark cycle. An overview of the experiment is provided in **Fig. S1**.

### Measurement of CT_Max_

To determine whether flies from different climates differed in their thermal tolerance, we measured the critical thermal maximum (CT_Max_), here defined as the temperature at which flies enter a heat coma, using thermal ramping experiments. Over a two-day period, 64 F2 female flies (32 from each population, aged 10-18 days after adult eclosion) were treated. Flies were transferred to 5ml tubes by aspiration, which were then sealed and inserted into a tube holder that was fully submerged in a water bath (PolyScience) at 25°C. The temperature was increased by 0.5°C every minute for 20 minutes (up to 35°C), then in 0.1°C increments every minute until all flies had passed out. When flies appeared to faint, tubes were tapped 3 times. Flies that got back up were kept in the water bath, while unresponsive flies were removed and transferred to an individual food tube to check for survival. Measurements were made on 16 flies at a time.

### F3 heat shock experiment

To assess the direct impacts of heat shock on chromatin accessibility and gene expression in the ovaries of F3 females, groups of females were either exposed to a heat shock (HS) of 37°C or ambient control (Ctrl) by immersion in one of two water baths for one hour, one heated to 37 °C, and the other unheated at 19-21°C. Ovaries were immediately dissected for use in ATAC-seq and RNA-seq experiments following this treatment. DNA-seq samples were prepared from the remaining tissues of control animals following ATAC-seq dissections.

For ATAC-seq experiments, 288 F3 female flies were treated (2 treatments, 2 populations, 3 replicates, each replicate a pool of 24 flies). HS and Ctrl treatments were applied to groups of 12 flies at a time, to reduce the time between treatment and dissection to less than 20 minutes. Ovaries were dissected in phosphate-buffered saline (PBS) and transferred to a microtube containing 200 µl PBS kept on ice. Once all individuals had been dissected, ovaries within replicates were homogenised using a dounce homogenizer (30 passes), and cells were counted using a Neubauer slide. Based on the average of five counts, 200,000 cells were isolated; these were centrifuged for 5 mins at 4°C, and then washed once with PBS at 4°C and resuspended in a freshly made lysis buffer (10 mM Tris·Cl, pH 7.4, 10 mM NaCl, 3 mM MgCl2, 0.1% (v/v) Igepal CA-630). The samples were incubated for 10 mins at 4°C, then centrifuged and the supernatant removed. These nuclei preparations were immediately transferred on ice to the sequencing facility for tagmentation and library preparation. The remaining tissues from Ctrl animals were then collected for DNA samples. Pooled samples of 24 females were collected in a microtube, spun down, and stored at -80°C. DNA extractions were carried out at a later date using the MagAttract HMW kit (Qiagen) following manufacturer’s instructions.

For the RNA-seq experiment, another set of 288 F3 female flies were treated in the same way as for ATAC-seq experiments. Once 12 ovary pairs (half a sample) had been dissected and added to a microtube containing 200 µl PBS, the microtube was briefly centrifuged, the supernatant was removed, and the sample was flash frozen in liquid nitrogen. Samples were then kept at -80°C, and extractions were carried out at a later date using the GenElute Mammalian Total RNA Miniprep kit (Sigma). The two halves of each sample were combined following homogenization in lysis buffer. The extraction was carried out according to manufacturer’s instructions, followed by an additional DNase I (Thermo Scientific) treatment and then precipitation with 5M lithium chloride and absolute ice cold ethanol.

Finally, the effects of F3 HS on F4 offspring development were measured in a phenotypic development assay of the offspring from 100 F3 females (2 treatments, 2 populations, 25 replicate lines). F3 virgin flies were collected and kept in single-sex food tubes at a density of 8-10 individuals. Five to seven day old virgin females were then exposed to HS and Ctrl treatments as described previously. Replicate lines were set up by transferring a single virgin female and 2 untreated virgin males from the same population to a new food tube (F4-1). Adults were transferred into a fresh food tube after two (F4-2), four (F4-3), six (F4-nonexperimental) and 12 (F4-4) days, and were removed from the experiment after 14 days. The following measurements were made for eggs laid in a two-day period in tubes F4-1,−2, -3, and -4: egg, pupae, and adult numbers; egg-to-pupa and egg-to-adult viability; and time to pupation and time to eclosion for eggs.

### Transgenerational effects of heat shock

To assess the transgenerational consequences of heat shock we repeated the RNA-seq and ATAC-seq experiments in the F6 and developmental phenotypic assays in the F7 (three generations after the original experiments). All flies used in transgenerational experiments were descendants of flies used in the F4 phenotypic development assay. Transgenerational experiments did not subject the flies to any further heat shock treatment but measured whether the consequences of heat shock in the F3 could still be detected in their descendants (dHS) compared with the descendants of controls (dCtrl).

ATAC-seq and RNA-seq samples in the F6 were carried out in the same manner as in the F3, except that in the preparation of ATAC-seq samples in the F6, fewer cells (100,000) were isolated than in the F3. Similarly, phenotypic development assays in the F7 were carried out in the same manner as in the F4, although only 18 out of the 25 lines from each treatment/population combination set up for the F4 phenotypic development assay were used.

### Statistical analysis of phenotypic traits

All statistical analysis of phenotypic traits was carried out in R (R Development Core Team 2022). CT_Max_ was analysed using the lmer function from the lme4 package (Bates 2010), with temperature as the response variable, population as a fixed effect and experimental batch as a random effect. Counts of eggs, pupae and adults were analysed with generalized linear models (glm) or generalized linear mixed effects models (the glmer function in lme4) with a poisson distribution. Egg-to-pupa and egg-to-adult viability data were also analysed using glm or glmer, but with a binomial distribution. Finally, age at pupation and age at eclosion were analysed with linear mixed effect models (lmer) with maternal ID as a random effect and the number of pupae in a tube considered as a covariate to account for differences in population density. Simple versions of models were carried out for analyses of F3 cohort 1, and mixed effects versions of models were used to analyse F3 cohorts 2-4 and F7 cohorts 1-4 (with cohort included as a random effect).

In the F4 and F7 phenotypic assays, heat shock treatment, population, and the interaction of the two factors were considered as fixed effects in statistical models. The significance of the interaction was tested using the R function dropterm (Venables and Ripley 2002), and if non-significant, the model was re-run with main-effects only. Chi-square or F statistics and P-values were obtained using the Anova function in the R package car (Fox and Weisberg 2019). If the interaction between treatment and population was significant, post-hoc tests were carried out using the emmeans package (Lenth 2020).

### Sequencing, bioinformatics and analysis of omics data

Library preparation and sequencing of DNA-seq, ATAC-seq, and RNA-seq samples was carried out by the CRG (Centre for Genomic Regulation, Barcelona Biomedical Research Park, Barcelona) Genomics Unit (including tagmentation of ATAC-seq samples). Samples were sequenced on an Illumina NextSeq 2000. A total of 6 DNA-seq libraries (Nextera DNA kit, 150 bp paired end reads), 24 ATAC-seq libraries (Nextera DNA kit, 50 bp paired end reads) and 24 RNA-seq libraries (Truseq Stranded mRNA with poly-A selection, 50 bp paired-end) were sequenced during the course of the experiment. An average of 25.2 million 150 bp PE reads (42 X coverage) were sequenced per sample for 6 DNA samples. Per sample average reads totaled, respectively, 48.2 million and 45.8 million for ATAC-seq and RNA-seq experiments in the F3, and 65.0 million and 35.6 million for ATAC-seq and RNA-seq experiments in the F6 (12 samples in 4 experiments, 48 total). All sequencing data has been deposited into the NCBI Sequence Read Archive under accession number PRJNA1002872.

We used three different tools to identify TE insertions in DNA samples. To find probable reference TEs matching those in release 6 version 46 of the *D. melanogaster* reference genome (Dmel_r6v46), we ran Tlex-3 (Bogaerts-Márquez et al. 2020) using a list of 2,417 reference TEs identified within the euchromatic region of the genome (Rech et al. 2022). TEs that were identified as present or polymorphic in all three samples from each population were included as ‘reference TEs’. To identify putative ‘non-reference TEs’ we used the consensus insertions identified by two different tools: PoPoolationTE2 (Kofler et al. 2016) and TEMP2 (Yu et al. 2021). Results for different software, samples, and populations were combined using the merge and intersect functions of bedtools (Quinlan and Hall 2010), with merge distance set to 25 bp. Intersecting insertions were only retained if the TEs’ families matched. For each population we included non-reference TE insertions that were present in that population in at least 2 out of 3 replicate DNA samples according to both programs. A list of reference and non-reference TE insertions within 1kb of annotated genes was created for each population using the bedtools window function.

Transcript abundance of RNA-seq samples was estimated using Kallisto (Bray et al. 2016) and an index of Dmel_r6v46 transcripts. Differential expression analyses were then carried out in R using the DESeq2 package (Love et al. 2014). Transcript counts were imported into R and transformed into gene-level abundance estimates using the tximport package (Soneson et al. 2015) before analysis with DESeq2. Six separate pairwise comparisons were carried out: (1) HS vs Ctrl, F3 Akaa, (2) HS vs Ctrl, F3 Manz, (3) dHS vs dCtrl, F6 Akaa, (4) dHS vs dCtrl, F6 Manz, (5) F3 Ctrl vs F6 dCtrl, Akaa, and (6) F3 Ctrl vs F6 dCtrl, Manz. For each pairwise comparison, an initial filtration excluded transcripts with a total count of less than 10, and which appeared in less than three out of the six samples. We also used the lfcshrink function to generate more accurate effect sizes, and an adjusted *P*-value of 0.05.

We focussed on differentially expressed genes (DEGs) in response to heat shock or ancestral heat shock (pairwise comparisons 1, 2, 3 and 4 described above), and then explored similarities and differences in responses to heat shock or ancestral heat shock through set analysis, i.e. overlapping DEGs between (A) F3 Akaa and F3 Manz, (B) F6 Akaa and F6 Manz, (C) F3 Akaa and F6 Akaa, and (D) F3 Manz and F6 Manz. For the latter two, we excluded DEGs that differed between generations in controls (i.e. those present in pairwise comparisons 5 and 6 described above). Overlapping sets of heat shock responsive DEGs were visualized with the R package eulerr (Larsson 2022).

Chromatin accessibility was analysed using the nextflow core pipeline nf-core/atac version 2.0 (Ewels et al. 2020; Patel et al. 2022), with bwa used to align reads to Dmel_r6v46. The nf-core/atac pipeline aligns and filters sequences, and calls per-sample ATAC-seq peaks relative to a merged consensus peak set. All 24 ATAC-seq samples (triplicated samples for 2 treatments, 2 populations and 2 generations) were considered to generate the consensus peak set. Sequence alignment map files (BAM) were then integrated with Kallisto transcript counts using the R package intePareto (Cao et al. 2020). Matching and integration steps of intePareto were run for the same six pairwise comparisons described for DESeq2 analysis of RNA-seq data, with patterns of transcription matched to the weighted mean values of promoter peaks (up to 1kb from the gene).

Once RNA-seq data and ATAC-seq datasets were integrated, we carried out differential accessibility analysis of the ATAC-seq data with DESeq2 (analysis of the same pairwise comparisons and using the same filtration parameters, treatment with lfcshrink, and *P*-values as described for the RNA-seq data). We then compared lists of genes with differentially accessible promoter regions (DARs) using set analysis in the same way as described for lists of DEGs. Differential expression/accessibility results and log fold change (LFC) values for all genes from both populations and both generations were then merged and combined with lists of TE insertions within 1kb of annotated genes, allowing us to investigate associations between expression, accessibility and the presence of TEs in both generations and populations.

The relationship between gene expression and accessibility in the F3 of each population was assessed using linear regression. Expression LFC was considered as the response variable and accessibility LFC as the explanatory variable. Linear models also included differential expression class (not DE, shared across populations, unique) as a term in the model and its interaction with DAR LFC. Linear regression was also used to investigate whether expression LFC in the F3 was predictive of expression LFC in the F6.

To look for associations between classes of TE insertions (no TE, reference TE or non-reference TE) and patterns of gene expression and chromatin accessibility in the F3, we carried out Chi-squared tests considering both whether genes were DE or DA, and the direction of change. In total we looked for associations between TEs and five expression/accessibility characteristics: i) differential expression class (not DE, shared, unique); ii) differential accessibility class (not DA, shared, unique); iii) direction of differential gene expression change; iv) direction of differential chromatin accessibility change, and; v) direction of expression and accessibility change across all genes (not just those with significant differences). Differences were confirmed using post-hoc tests from the chisq.posthoc.test package (Ebbert 2019).

We tested for functional enrichment of GO biological processes for each population using the enrichGO function of the clusterProfiler package (Wu et al. 2021), simplifying lists of GO terms with the rrvgo package (Sayols 2020). We carried out enrichment for the following: i) genes that were both DEGs and DARs in the F3; ii) genes that were DEGs in both the F3 and F6; and iii) genes associated with TEs that were positively associated with directional changes in DEGs or DARs. In the first two cases, genes were split into four groups based on the direction (positive or negative) and concordance (concordant or discordant) of change between F3 expression and F3 accessibility, or F3 expression and F6 expression.

## Acknowledgements

We would like to thank Miriam Merenciano for her help with experimental dissections and Marta Coronado-Zamora for assistance and discussion about bioinformatic analyses. We also thank Simon Orozco Arias and María Bogaerts-Márquez for their helpful bioinformatic advice. Flies from Akaa, Finland, were collected by Maaria Kankare from the University of Jyväskylä, and flies from Manzanares, Spain, were collected by secondary school teachers and pupils from IES (Instituto de Enseñanza Secundaria) Azuer as part of the citizen science project Melanogaster Catch the Fly!

E.H. received funding from the European Union’s Horizon 2020 research and innovation programme under the Marie Sklodowska-Curie grant agreement No 101030460, and J.G. was supported by Grant PID2020-115874GB-I00 funded by the Spanish Ministry of Sciences and Innovation MCIN/ AEI /10.13039/501100011033, by grant PID2023-148838NB-I00 funded by the MCIU/AEI/10.13039/501100011033/FEDER, EU, and by grant 2021 SGR 00417 funded by Departament de Recerca i Universitats, Generalitat de Catalunya. The citizen science project was supported by grant FCT-20-15710 funded by FECYT-Ministerio de Ciencia, Innovación y Universidades.

## Data Availability

Genomic data are available from the NCBI (https://www.ncbi.nlm.nih.gov/) Sequence Read Archive under accession number PRJNA1002872. Phenotypic data, processed sequence data and the scripts that were used to generate the findings of this study are openly available from Digital.CSIC (https://digital.csic.es/) under DOI https://doi.org/10.20350/DIGITALCSIC/17069.

## Notes

https://www.ncbi.nlm.nih.gov/bioproject/PRJNA1002872/

### Competing Interest Statement

The authors have declared no competing interest.

