## Supplementary Figures for "Transgenerational effects of heat shock on gene regulation and fitness-related traits are stronger in arid than temperate *Drosophila* populations"

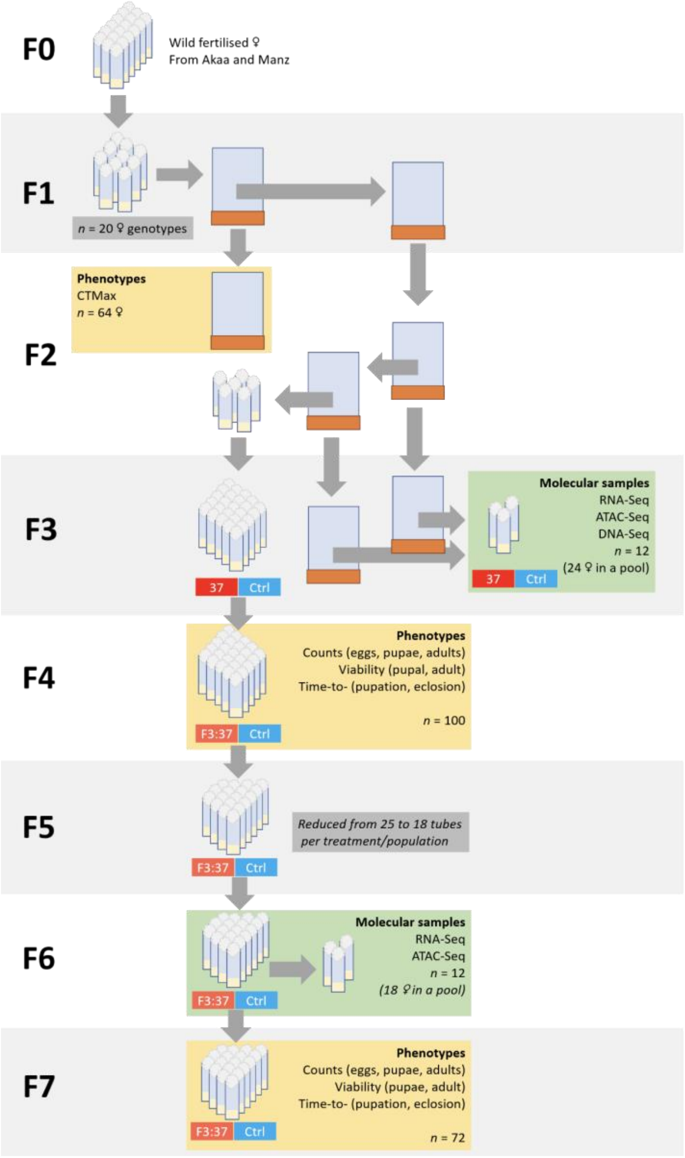


***Fig.S1.*** *Schema of multigeneration phenotypic and molecular experiments from establishment of F0 lines through to phenotypic measures taken in the F7. Heat shock treatment was applied in the F3, subsequent experiments considered the transgenerational effects (without additional treatment). Sample sizes (n) refer to total replicates in molecular experiments (all treatments and populations), and number of tubes in phenotypic studies. In the F4 and F7 multiple measures were made in each tube, but 'tube' was always considered as a random effect in statistical models. Populations were treated in the same way throughout the experiment.*

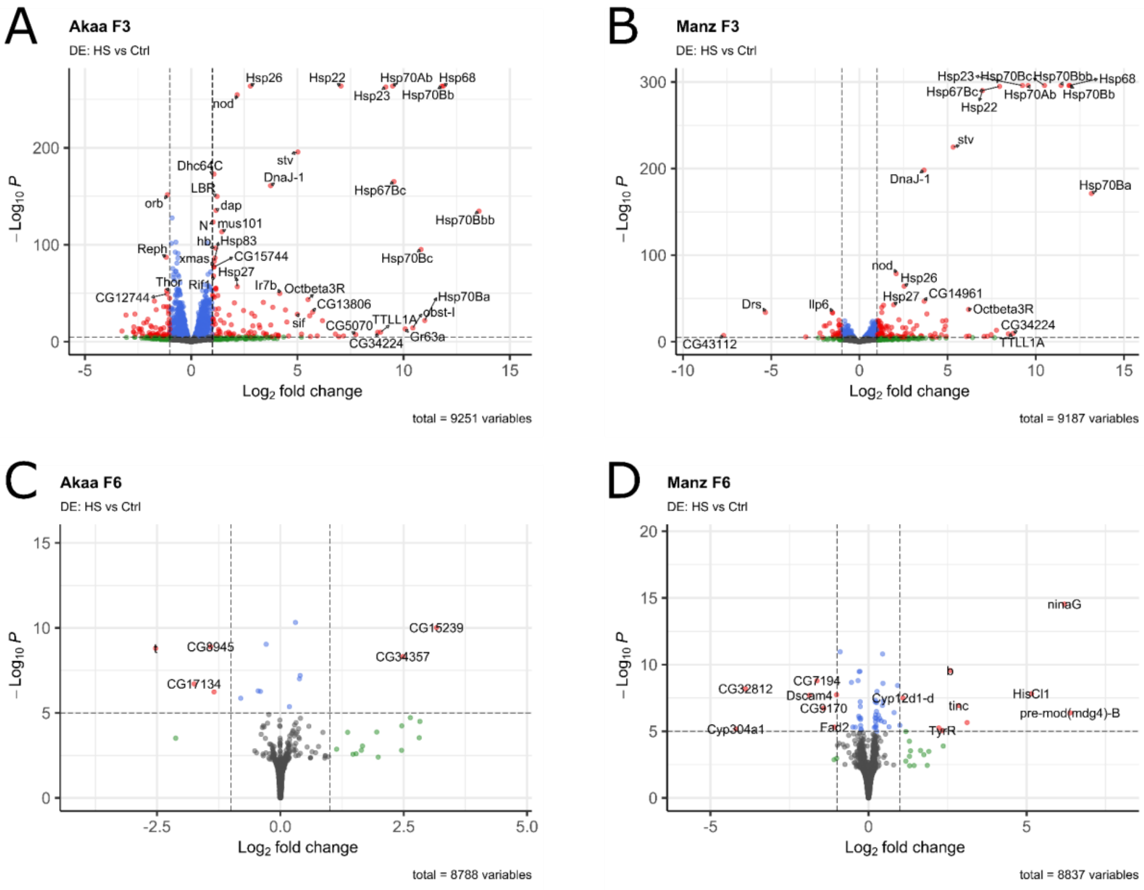


***Fig. S2.*** *Volcano plots show the effect of F3 heat shock of gene expression in (A) F3 Akaa ovaries, (B) F3 Manz ovaries, (C) F6 Akaa ovaries, and (D) F6 Manz ovaries, with some highly significant genes marked. For all four plots, an FDR corrected P-value of 0.05 is used. In (A) and (B), an L2FC of 1 is used to highlight larger effect sizes (equivalent to a fold change of 2). In (C) and (D), an L2FC of 0.585 is used to highlight larger effect sizes (equivalent to a fold change of 1.5).*

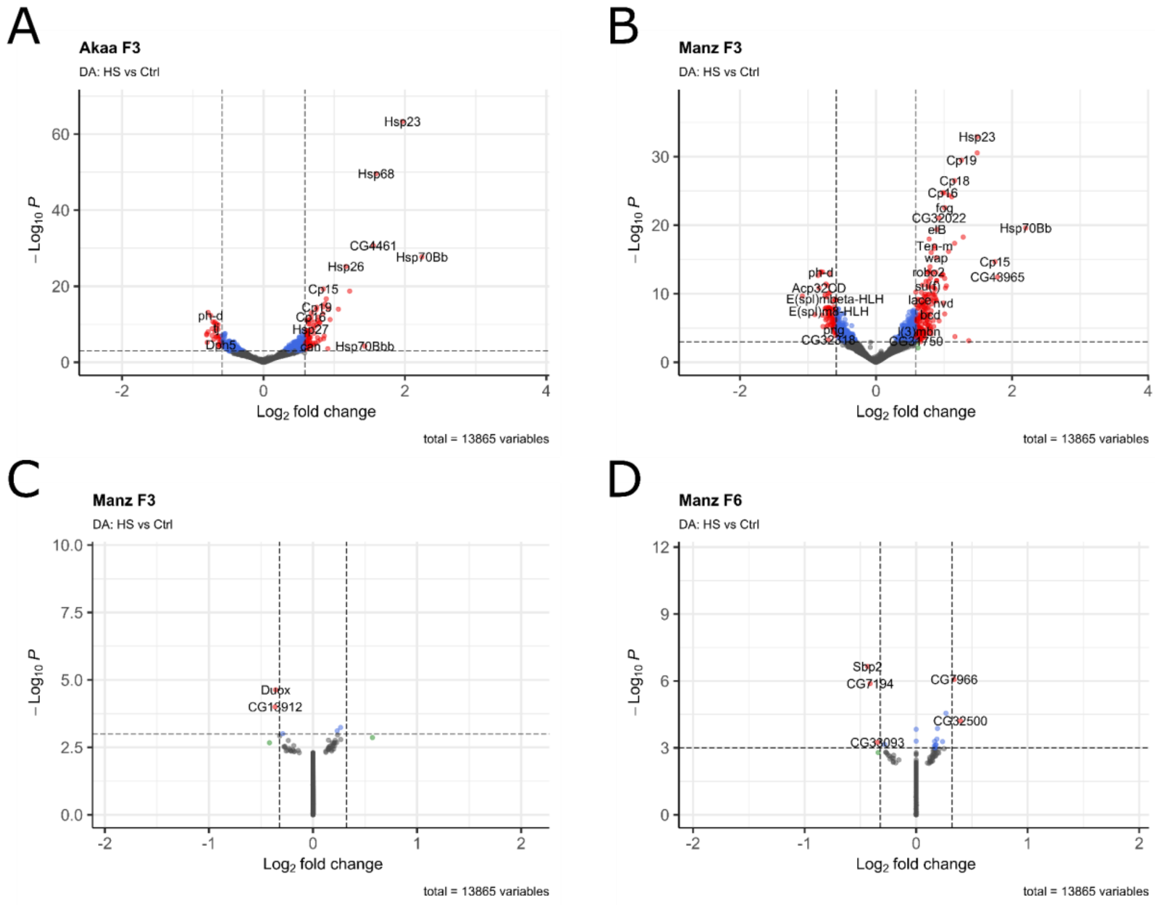


***Fig. S3.*** *Volcano plots show the effect of F3 heat shock of chromatin accessibility in (A) F3 Akaa ovaries, (B) F3 Manz ovaries, (C) F6 Akaa ovaries, and (D) F6 Manz ovaries, with some highly significant genes marked. For all four plots, an FDR corrected P-value of 0.05 is used. In (A) and (B), an L2FC of 0.585 is used to highlight larger effect sizes (equivalent to a fold change of 1.5). In (C) and (D), an L2FC of 0.3219 is used to highlight larger effect sizes (equivalent to a fold change of 1.25).*

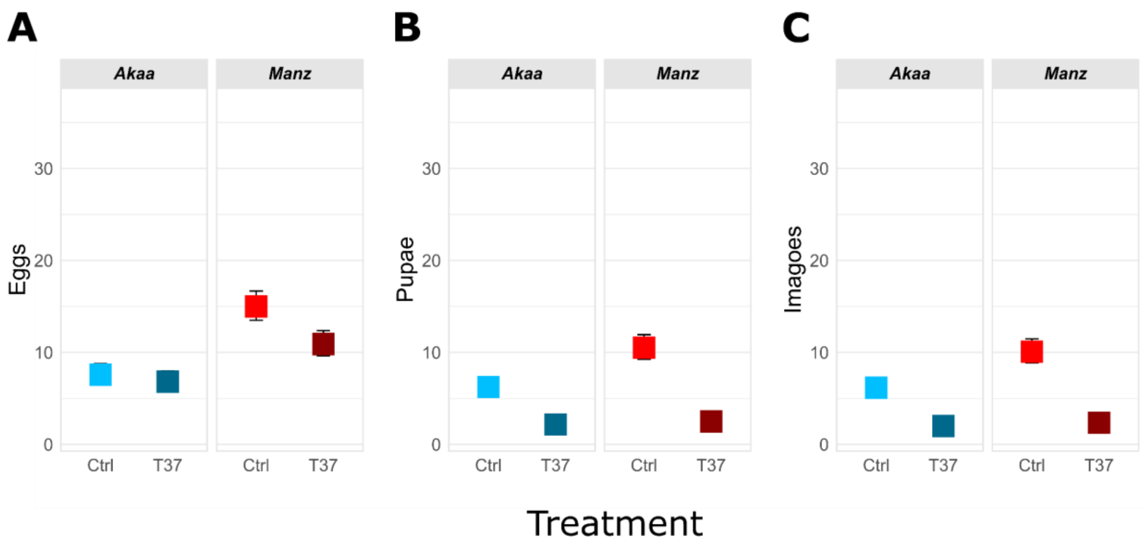


***Fig. S4.*** *Effect of heat shock (T37) on absolute numbers of (A) eggs, (B) pupae, and (C) adults for offspring produced within 48 hours of treatment (cohort 1). Y-axes are kept constant in all panels of Figs. S1 and S2 to allow comparison across life stages and cohorts.*

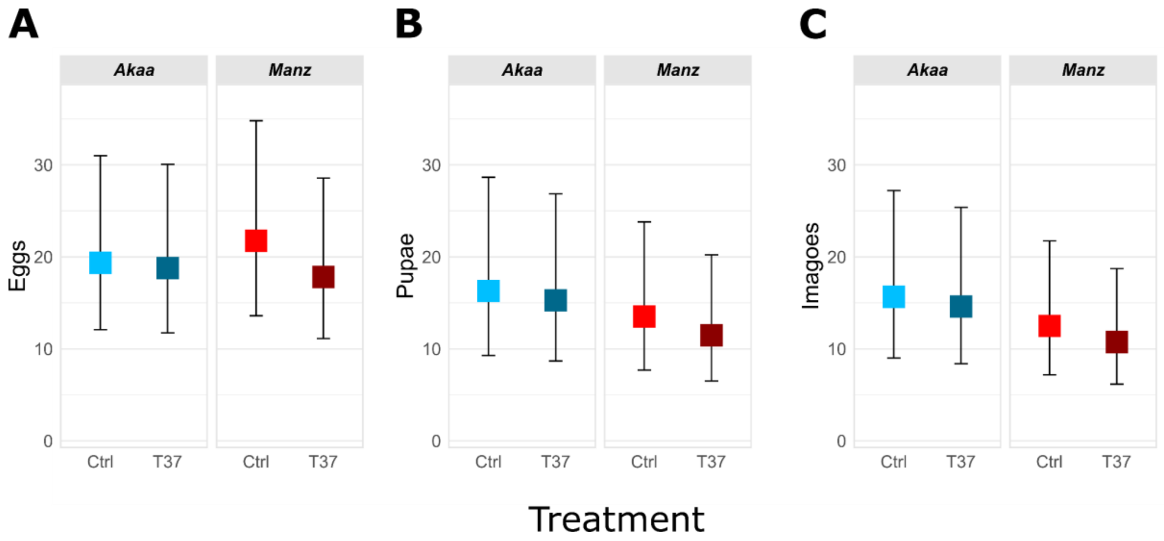


***Fig. S5.*** *Effect of heat shock (T37) on absolute numbers of (A) eggs, (B) pupae, and (C) adults for offspring produced in 3 separate 48 hour periods between 2 and 14 days after treatment (cohorts 2-4). Y-axes are kept constant in all panels of Figs. S1 and S2 to allow comparison across life stages and cohorts.*

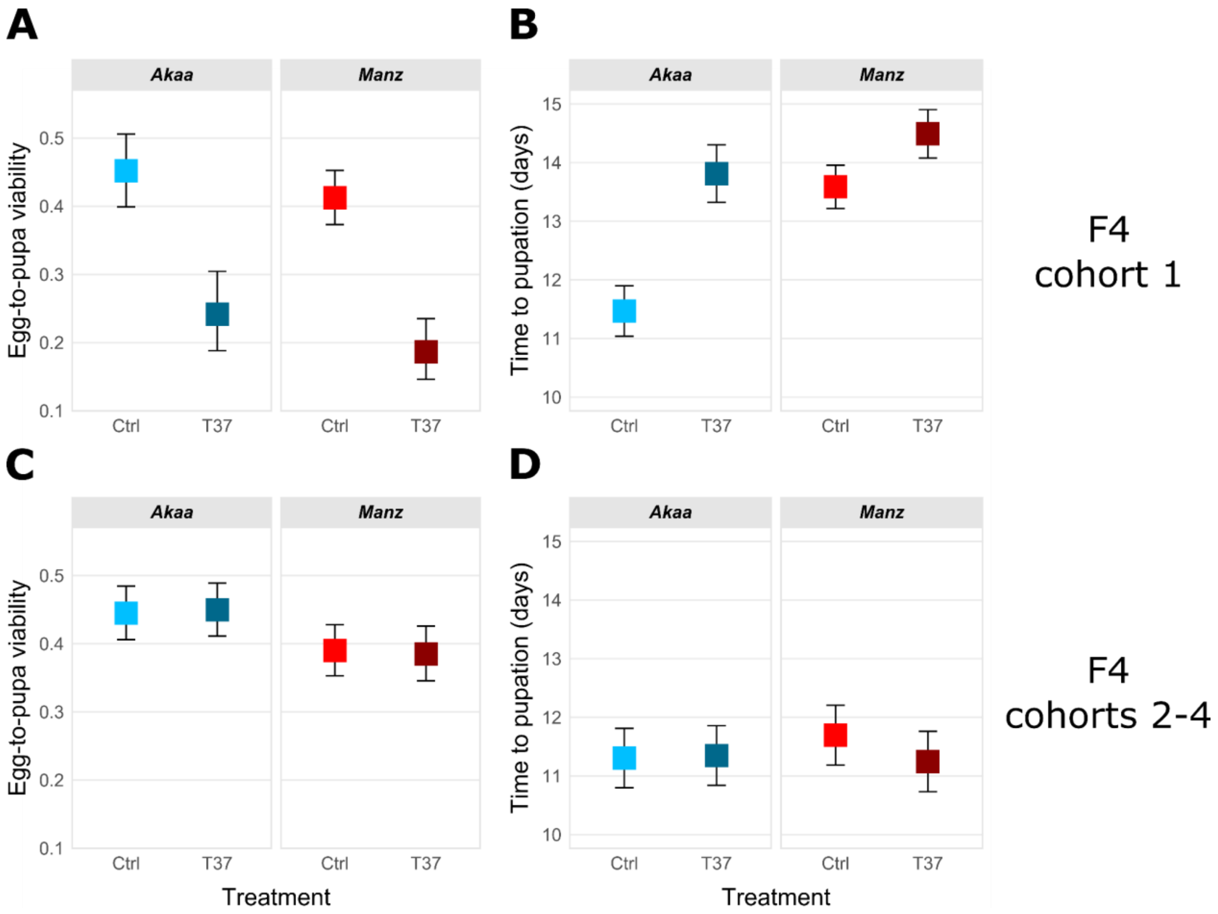


***Fig. S6.*** *Effect of heat shock (T37) on F4 offspring (A) egg-to-pupa viability and (B) time to pupation in cohort 1 and (C) egg-to-pupa viability and (D) time to pupation in cohorts 2-4.*

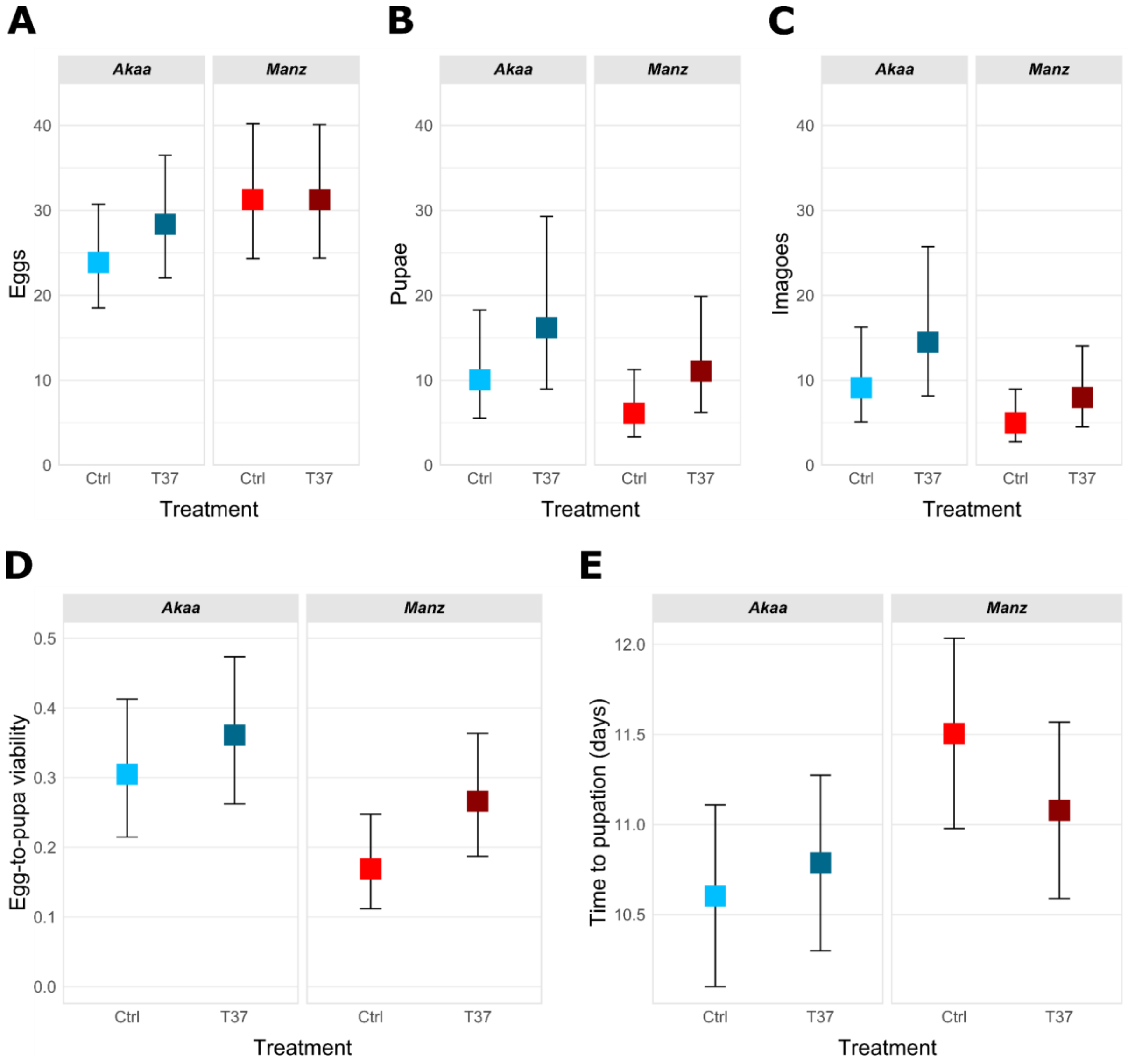


***Fig. S7.*** *Effect of ancestral heat shock (T37) on F7 offspring absolute numbers of (A) eggs, (B) pupae, and (C) adults, and on (D) egg-to-pupa viability, and (E) time to pupation of great-grand offspring from all four cohorts measured in the F7 generation.*
